# Stony coral populations are more sensitive to changes in vital rates in disturbed environments

**DOI:** 10.1101/2020.02.17.952424

**Authors:** Tessa E. Hall, Andrew S. Freedman, André M. de Roos, Peter J. Edmunds, Robert C. Carpenter, Kevin Gross

## Abstract

Reef-building corals, like many long-lived organisms, experience environmental change as a combination of separate but concurrent processes, some of which are gradual yet long-lasting, while others are more acute but short-lived. For corals, some chronic environmental stressors, such as rising temperature and ocean acidification, are thought to induce gradual changes in colonies’ vital rates. Meanwhile, other environmental changes, such as the intensification of tropical cyclones, change the disturbance regime that corals experience. Here, we use a physiologically structured population model to explore how chronic environmental stressors that impact the vital rates of individual coral colonies interact with the intensity and magnitude of disturbance to affect coral population dynamics and cover. We find that when disturbances are relatively benign, intraspecific density dependence driven by space competition partially buffers coral populations against gradual changes in vital rates. However, the impact of chronic stressors is amplified in more highly disturbed environments, because disturbance weakens the buffering effect of space competition. We also show that coral cover is more sensitive to changes in colony growth and mortality than to external recruitment, at least in non-self-seeding populations, and that space competition and size structure mediate the extent and pace of coral-population recovery following a large-scale mortality event. Understanding the complex interplay among chronic environmental stressors, mass-mortality events, and population size structure sharpens our ability to project coral dynamics in an increasingly disturbed future.

## Introduction

Tropical reef-forming corals are celebrated worldwide for the spectacular and diverse marine communities they support (Huston, 1985; Knowlton et al., 2010), and for the ecosystem services that those communities provide (Moberg and Folke, 1999; Knowlton et al., 2010; Wild et al., 2011). Reef-building corals are also thought to be particularly vulnerable to ongoing environmental changes (Hoegh-Guldberg et al., 2007; Hughes et al., 2017), including rising sea temperatures, acidifying oceans, increased competition from macroalgae, and increased nutrient run-off and sediment loading in coastal waters. The aggregate effect of these changes may place many reefs in jeopardy of substantial ecological restructuring, if not wholesale collapse.

A particular challenge to projecting how reefs will respond to environmental change is that different components of environmental change will have different demographic consequences. In the case of corals, many coral reefs intermittently experience catastrophic mortality events from disturbances such as typhoons, predator outbreaks, bleaching, and disease. Environmental change is expected to increase the frequency and/or intensity of these disturbances (for example, tropical storms have recently become more powerful (Emanuel, 2005)). However, other aspects of environmental change — most notably rising sea temperatures and ocean acidification (OA) — simultaneously present chronic stressors that will alter the physiology and vital rates of individual coral colonies. For example, increasing temperature or decreasing calcification rates may reduce growth rates (De’ath et al., 2009); reduced skeletal density and/or increased microbioerosion might make colonies more susceptible to fragmentation or dislodgement from hydrodynamic stress (Madin et al., 2012; Reyes-Nivia et al., 2013); and recruitment of new colonies may be reduced by sublethal effects of bleaching (Ward et al., 2002) or increased macroalgal cover (Kuffner et al., 2006). Forecasting the aggregate impact of environmental change requires understanding how physiological responses to chronic stressors interact with changes in the disturbance regime that corals experience. For example, Ortiz et al. (2018) recently showed that recovery rates from disturbances have slowed on the Great Barrier Reef, and suggested that these slower recovery rates may be a consequence of the combined effects of several chronic stressors.

Yet the interaction between chronic stressors and acute disturbances may be more nuanced than simply repeated episodes of death and recovery. In particular, size structure is known to be an important mediator of coral population dynamics. Over 30 years ago, Roughgarden et al. (1985) showed that sessile marine invertebrates with long-distance larval dispersal can undergo sustained population cycles driven by the interaction between age structure and density-dependent recruitment. Even when population cycles do not persist in perpetuity, populations may show damped oscillations in the recovery phase following a catastrophic disturbance. Pascual and Caswell (1991) showed that the same oscillations can appear in size-structured populations, thus establishing an important connection to stony corals, as vital rates of coral colonies often depend on colony size (Hughes, 1984; Hughes and Connell, 1987). Later work by Artzy-Randrup et al. (2007) suggested that density-dependent growth can stabilize these general dynamics and thus make population cycles less likely. Nevertheless, this theory makes it clear that a full consideration of coral populations’ response to environmental change must account for the demographic consequences of size structure within the coral population.

The objective of this study is to investigate how gradual changes in coral colonies’ vital rates driven by chronic stressors will interact with an increase in the frequency and intensity of disturbances, when vital rates depend on both colony size and population density. We investigate this question by developing a physiologically structured population model (PSPM) of stony corals, building on theory developed by de Roos and colleagues (de Roos, 1997; Kirkilionis et al., 2001; de Roos et al., 2010; de Roos and Persson, 2013). Our core analysis consists of two parts. First, we investigate how size structure impacts coral population dynamics when coral populations experience stochastic mortality pulses from disturbances such as typhoons or predator outbreaks. Second, we conduct an elasticity analysis to quantify how changes in vital rates driven by chronic stressors might scale up to affect average coral cover, and ask how these sensitivities may depend on the frequency and intensity of mortality pulses.

Stony coral themselves are a hugely diverse taxon, with a vast array of growth morphologies, reproductive modes, and habitats. We do not attempt to capture the full range of this diversity here. Instead, we parameterize our model using data for the *Pocillopora verrucosa* species complex (see Edmunds et al. (2016)) as it occurs at 10 m depth on the fore reef of the north shore of Mo’orea, French Polynesia. We choose this species complex and habitat because it is well-studied and common on Mo’orea (Comeau et al., 2016; Holbrook et al., 2018; Doo et al., 2019), and has been a focus of other modeling efforts (Kayal et al., 2018). Our modeling framework is sufficiently general that it can be customized to other coral populations, although comparisons among different coral species are beyond the scope of this article.

The rest of this paper is structured as follows. In the Methods section, we introduce the mathematical model and describe how it is parameterized for *P. verrucosa*. Our analysis of the model, presented in the Results section, proceeds in two parts. First, we investigate the population dynamics predicted by the model under both undisturbed and intermittently disturbed conditions. Second, we conduct an elasticity analysis to investigate how coral cover responds to changes in colonies’ vital rates under different disturbance regimes. Technical details about parameter estimation and model analysis are provided in the supplement.

## Methods

### Mathematical model

Our development closely follows de Roos (1997). We classify individual coral colonies by their size, because size is often correlated with the demographic fate of individual colonies (Hughes and Connell, 1987). We measure a coral colony’s size by its effective diameter *x*. In other words, if a colony’s planar area is *A*, then the effective diameter *x* is defined by the familiar relation *A*(*x*) = *π* (*x*/2)^2^. Let *n*(*t, x*) give the density of coral colonies with size *x* at time *t*. Note that *n*(*t, x*) is a density (in the sense of a probability density) of a population density (in the sense of number of colonies per *m*^2^); that is, the population density of corals at time *t*, in terms of colonies per *m*^2^, is *n*(*t*) = *∫ n*(*t, x*) *dx*.

Let *x*_0_ be the size of a newly settled and metamorphosed planula (i.e., a single polyp, often referred to as a spat) and let *x*_max_ be the maximum achievable colony size in the local habitat. The total coral cover at time *t*, measured as the proportion of available substratum covered, is given by

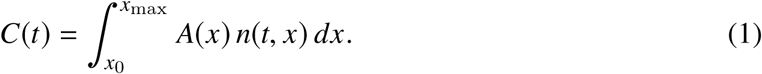

The proportion of available substratum is 1 − *C*(*t*). (As will be seen below, the model is defined to ensure 0 ≤ *C*(*t*) ≤ 1.) Although our definitions of *C*(*t*) and 1 − *C*(*t*) are most natural in the context of a monoculture, for a multi-species reef *C*(*t*) can be interpreted as the cover of the focal population as a proportion of the habitat available to that species.

Three vital rates govern the predicted population dynamics. Each vital rate may depend on both a colony’s size and the population’s total cover. The growth rate of a colony is the rate at which its size changes with respect to time, written as *g*(*x, C*). We assume that growth is deterministic, and depends only on the colony’s size and the total population cover. The second vital rate is the background mortality rate, written as *µ*(*x, C*). This gives the rate at which colonies perish from chronic mortality, such as dislodgement, overgrowth, or disease. This rate does not include episodic mass mortality caused by mass bleaching, cyclones, or predator outbreaks. For simplicity, we assume that when a colony dies, the space that it occupied immediately becomes available to other living colonies. We also do not consider partial mortality, which may be important for many coral species.

The third vital rate is the recruitment rate. Following previous theory (Roughgarden et al., 1985; Artzy-Randrup et al., 2007), we will consider a single coral population that recruits predominantly via larval immigration; internal recruitment (i.e., self-seeding) is assumed to be negligible. Successful recruitment is limited by the available free space (Hughes and Jackson, 1985; Connell et al., 1997). The rate at which new recruits arrive and settle is written as *s*(*C*).

Collecting all model components gives the full model for coral population dynamics as (de Roos, 1997)

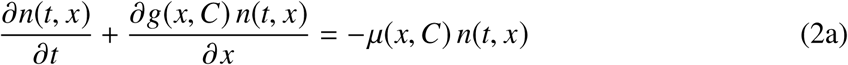

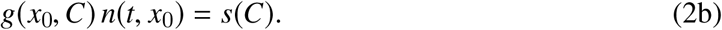

Eq. 2a is a balance equation that relates the change in the density of coral colonies to their growth and mortality. Eq. 2b is a boundary condition that adds new individuals to the population via external recruitment.

### Parameterization for *Pocillopora verrucosa*

We parameterize our model based on the species complex *Pocillopora verrucosa* as it is found at 10 m depth on the north shore of Mo’orea, French Polynesia (Edmunds et al., 2016). No single data set contains all of the information that we need to parameterize the model fully. Thus, we pool information from several sources, including previously published data (Holbrook et al., 2018), our own annual photoquadrat monitoring data, and precedent set in earlier modeling studies (Muko et al., 2001; Artzy-Randrup et al., 2007). Details about parameter estimation can be found in the supplement.

We assume that the maximum attainable size of a *P. verrucosa* colony is *x*_max_ = 50 cm (Veron, 2000), and that a newly settled coral polyp has a diameter of *x*_0_ = 0.4 mm (Babcock, 1991). To quantify growth rates, we first assume that the growth rate *g*(*x, C*) takes the form

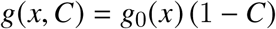

where *g*_0_(*x*) gives a colony’s growth rate in uncrowded conditions. Density-dependence is assumed to reduce growth in proportion to the available free space, equal to 1 − *C* (Muko et al., 2001; Artzy-Randrup et al., 2007). To estimate *g*_0_(*x*) we use annual photoquadrat monitoring data of individual *P. verrucosa* colonies from this habitat for 2011–2018. These years followed a large die-off caused by an outbreak of the crown-of-thorns seastar in 2002–10 and a cyclone in 2010 (Kayal et al., 2012; Holbrook et al., 2018). Thus, most coral colonies observed during these surveys were small (*x* ≤ 12 cm). These data suggest that as corals become larger, coral growth rates increase, but at a decelerating rate, at least across the size ranges found in this data set. Thus, we fit a quadratic curve for *g*_0_(*x*), constrained so that the growth rate is 0 when a colony attains its maximal size, i.e., *g*_0_(*x*_max_) = 0 (Fig. 1A).

**Figure 1:**
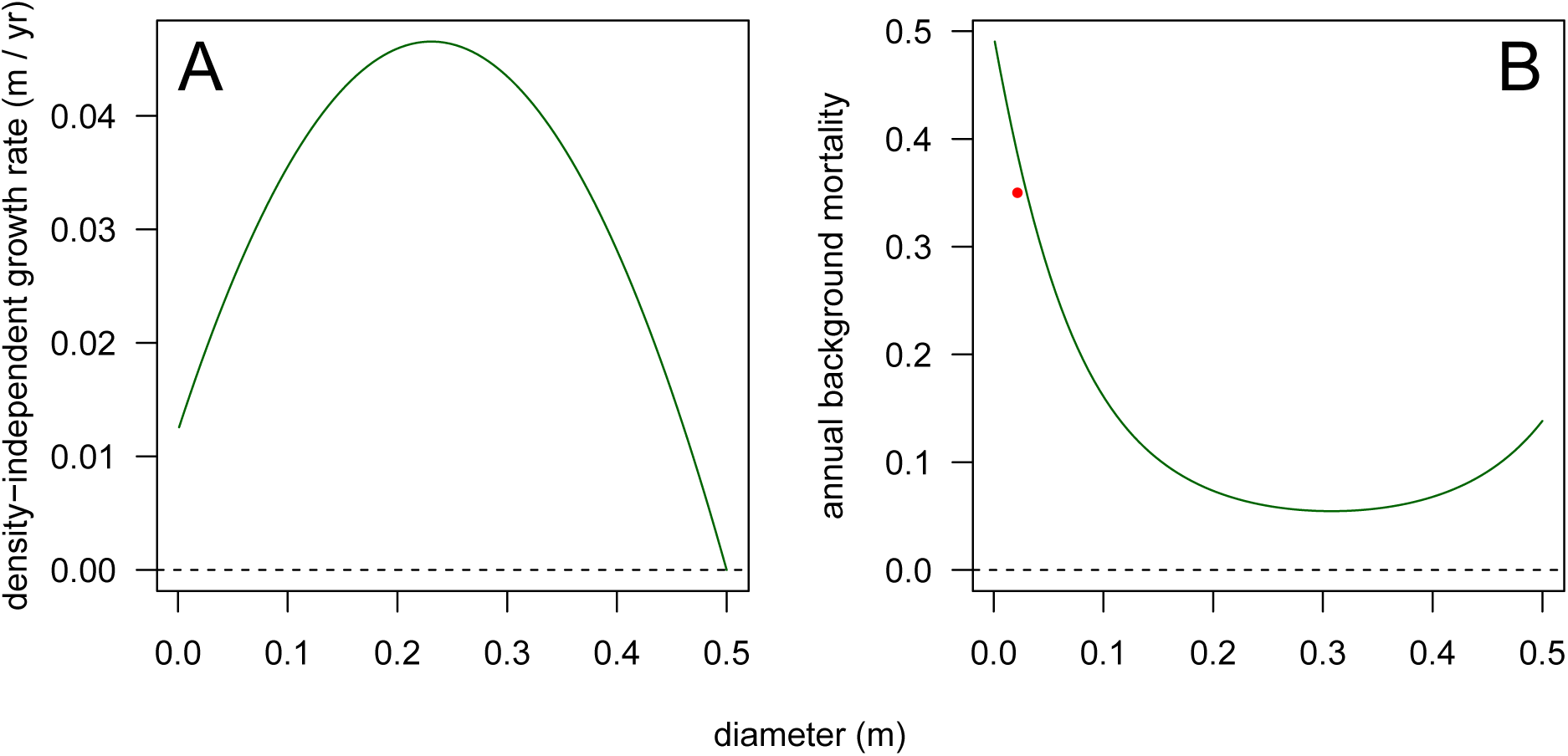
Growth and mortality functions used in the model. A: Density-independent growth rate in m / yr vs. colony diameter. This plot shows *g*_0_, the growth rate in uncrowded conditions. B: Annual mortality vs. colony diameter, based on data for corymbose colonies reported in Madin et al. (2014), and adjusted to match 35% small-colony mortality for *x* = 0.03 m colonies reported in Holbrook et al. (2018) (shown by the point in red). This plot shows annual mortality as a proportion, although mortality enters the model as the equivalent rate.

We assume that mortality is density-independent, thus *µ*(*x, C*) = *µ*(*x*). Following Madin et al. (2014), we expect that very small and very large colonies will experience larger mortality than intermediate-sized colonies; that is, the relationship between size and mortality will be “u-shaped”. This shape arises because small coral colonies are more vulnerable to a wide variety of mortality types including overgrowth from space competitors, whereas large colonies are more vulnerable to dislodgement from hydrodynamic stress (Madin et al., 2014). Our photoquadrat data do not include enough large colonies to estimate the full relationship between mortality and colony size. Thus, we instead use size-specific mortality data reported by Madin et al. (2014) for corymbose corals at Lizard Island, Australia. Of the coral growth forms included in Madin et al. (2014), the corymbose growth form is most similar to the closed branching growth form of *P. verrucosa*, and thus we expect the qualitative shape of the mortality curve to be similar. However, we also expect that the acroporid species used by Madin et al. (2014) will have higher mortality than *P. verrucosa*, as verified by Holbrook et al. (2018)’s mortality data for *Pocillopora* recruits at Mo’orea. Thus, we fit a size-dependent survival curve to Madin et al. (2014)’s corymbose colony data, and then multiplied this curve by a constant factor to adjust the small-colony mortality to match the mortality from Holbrook et al. (2018) (Fig. 1B).

Finally, we assume that recruits arrive at a baseline rate *s*_0_. Recruits successfully settle at a rate proportional to the amount of free space (Hughes and Jackson, 1985; Connell et al., 1997), giving

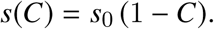

Holbrook et al. (2018) report that *P. verrucosa* recruits large enough to be easily detectable in photoquadrats (*x* = 0.03 m) are found on the north shore of Mo’orea at an average rate of approximately 20 m^−2^ y^−1^. Our growth and survival curves suggest that in uncrowded conditions roughly 40% of all recruits will survive long enough to grow to *x* = 0.03 m, giving an arrival rate of newly settling recruits of *s*_0_ ≈ 20/.4 = 50 recruits m^−2^ y^−1^.

## Results

First, we explore the population dynamics generated by the model under our baseline parameter set, both with and without mass-mortality events caused by disturbances such as typhoons or bleaching. Second, we conduct a sensitivity analysis to investigate how changes in vital rates driven by chronic stressors such as temperature or OA interact with mortality pulses to affect overall coral cover.

### Population dynamics

Figure 2A shows total coral cover over time for 100 (undisturbed) years following a catastrophe that completely eliminates the resident coral population. Recovery following a catastrophe is characterized by transient oscillations of considerable magnitude that eventually decay as coral cover approaches a stable equilibrium. These oscillations have a period on the order of one full oscillation per 40–50 years, suggesting that coral populations may exhibit slow dynamics that unfold on the time scale of scientific careers, or longer.

**Figure 2:**
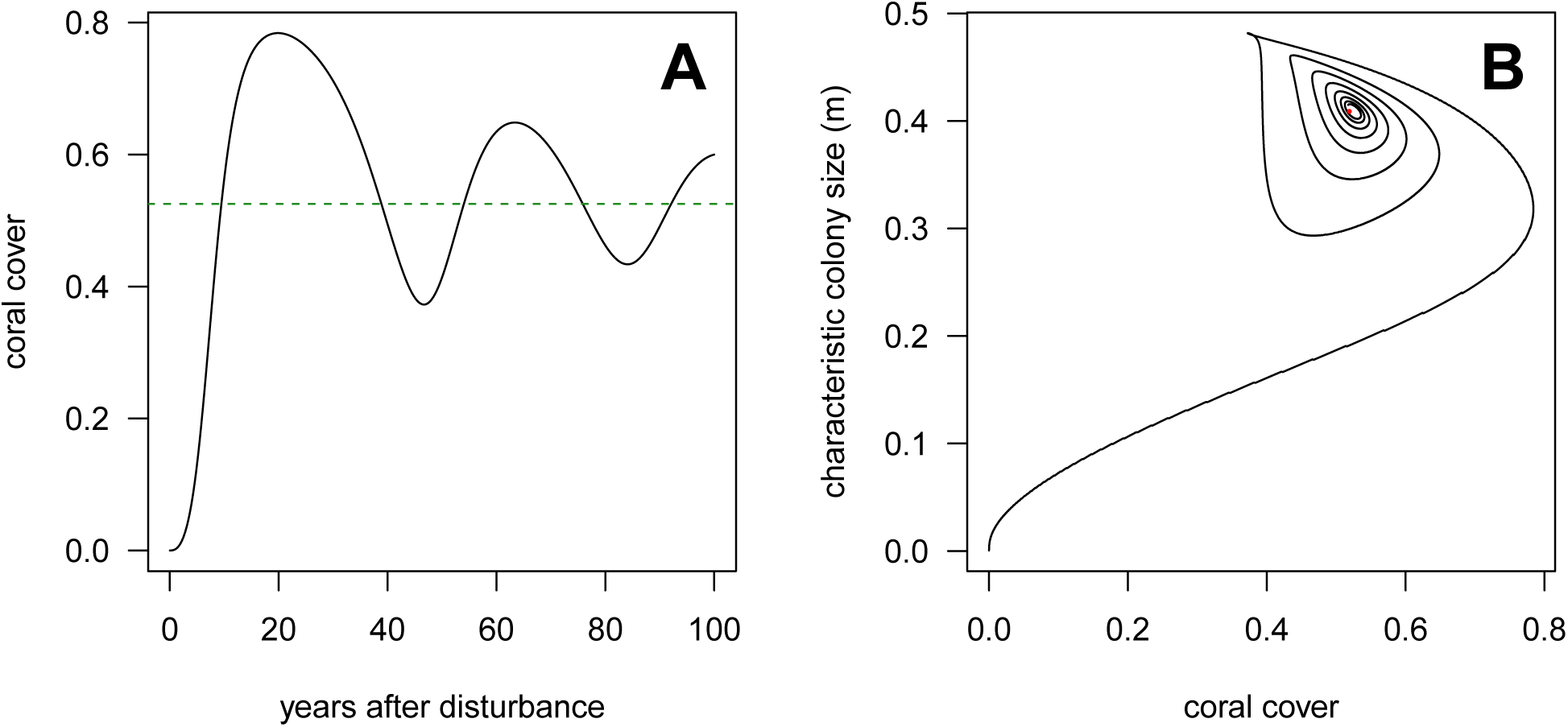
Model dynamics under the baseline parameter set for *P. verrucosa*. A: Total coral cover (as a proportion of available substrate) vs. time for 100 undisturbed years following a catastrophe that completely eliminates existing coral cover. The dashed horizontal line shows the equilibrium cover to which the dynamics eventually converge. B: Phase portrait showing the size (diameter) of a characteristic coral colony vs. the coral cover of the population. The characteristic colony size is defined such that colonies smaller than the characteristic size account for exactly half of the coral cover. Panel B shows 300 years of undisturbed dynamics following a catastrophe, and the red dot shows the equilibrium point to which the dynamics eventually converge.

These transient oscillations in coral cover are driven by space competition between cohorts of colonies (Figs. 2B, S1). Immediately after a catastrophe, the reef is repopulated by immigrating larvae from other reefs. These new recruits grow quickly in uncrowded conditions, rapidly occupying the available space. As cover builds, crowding reduces both growth and subsequent recruitment, leading to a population that consists mostly of corals that are large enough to escape overgrowth, yet small enough to avoid being dislodged. Eventually these colonies grow large enough that dislodgment mortality increases, opening new substratum to usher in the next cohort of recruits. These population cohorts echo the dynamics observed in early theoretical work for sessile marine invertebrates with space-limited recruitment (Roughgarden et al., 1985; Pascual and Caswell, 1991; Artzy-Randrup et al., 2007), and can be visualized by plotting coral cover against a characteristic colony size (Fig. 2B). Here, we define the characteristic colony size such that colonies smaller than the characteristic size account for half the coral cover, and colonies larger than the characteristic size account for the other half of coral cover. (We favor summarizing size structure in this way because the average or median colony size is affected by the size structure’s skew.) Fig. S1 provides an alternative visualization of these population oscillations.

For contemporary coral reefs, decades-long runs of undisturbed conditions seem unlikely. To ask if similar dynamics appear in disturbed environments, we conducted simulations with stochastic mass-mortality events. We modeled two disturbance regimes. In the moderate-disturbance regime (Fig. 3A,B), each year had a 10% chance of an event such as bleaching or a typhoon in which 25% of all colonies die (what we will subsequently call a “25% mortality event”), and a 4% chance of an 80% mortality event (such as a predator outbreak). In the high-disturbance regime (Fig. 3C,D), each year had a 20% chance of a 40% mortality event, and a 5% chance of a 95% mortality event. Mortality in these events was uniform across colony size.

**Figure 3:**
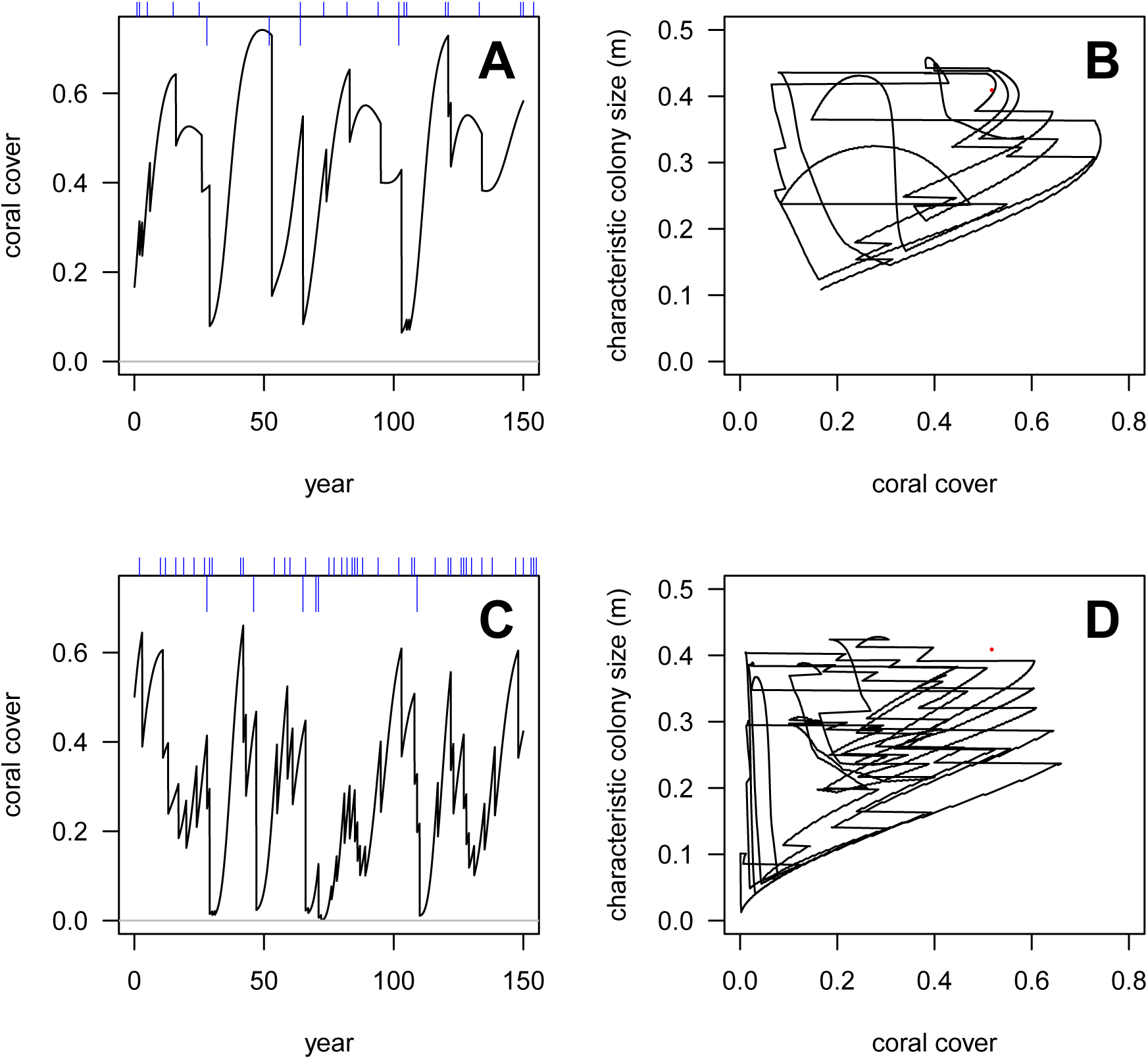
Population oscillations driven by size structure persist in moderately disturbed environments, but not in highly disturbed environments. A: Total coral cover (as a proportion of available substrate) vs. time for 150 years in the “moderate-disturbance” regime. Tick marks on the top axis show the timing of small (above the axis) or large (below the axis) mortality pulses. B: Characteristic colony size vs. coral cover for the same dynamics. Characteristic colony size is defined in the same way as in Fig. 2. The red dot shows the equilibrium under undisturbed conditions. C,D: Parallel plots for the “high-disturbance” regime, where mortality events are more frequent and more severe. Displayed dynamics follow a “burn-in” period of 200 years to eliminate transients.

These simulations show that size structure still mediates the recovery of coral populations following a widespread mortality event, even if fully formed population oscillations do not appear. Specifically, as long as a disturbance does not decimate a population (that is, when post-disturbance cover is ≥ 10%), the characteristic colony size will continue to increase following the disturbance (Fig. 3B,D). This suggests that, following a mild or moderate disturbance, recovery is initially driven by growth of surviving colonies as opposed to recruitment of new colonies. Consequently, cover will rebound more rapidly if the colonies that survive the disturbance are medium-size colonies that are able to grow most quickly in uncrowded conditions (Fig. 1A). Surviving colonies then grow rapidly to fill newly vacated space, to the extent that this growth is not impeded by dead coral skeletons. On the other hand, if the disturbance occurs when the population is dominated by large colonies, then physiological constraints will limit the survivors’ post-disturbance growth, despite the newly relaxed intraspecific competition. Thus, the population may fail to rebound to pre-disturbance cover, even if the years following the disturbance are benign. On the other hand, when a severe disturbance reduces the coral population to very low cover (≤ 10%), the surviving colonies are too sparsely distributed to rebuild cover through their growth. Instead, recovery must wait for a new cohort of recruits to arrive and to replenish the population.

### Elasticity analysis

We now investigate how changes in colony-level vital rates driven by chronic disturbances such as rising sea temperatures or OA affect population cover. To do so, we quantify how a proportional change in a vital rate translates into corresponding proportional change in total cover; in other words, we conduct an elasticity analysis (de Kroon et al., 1986). To simplify matters, we suppose that changes in growth, mortality, or recruitment act independently of both colony size and coral density. Mathematically, we introduce multiplication factors *ϕ*_*g*_, *ϕ*_*µ*_ and *ϕ*_*s*_, such that the modified growth, mortality, and recruitment rates are *ϕ*_*g*_ × *g*(*x, C*), *ϕ*_*µ*_ × *µ*(*x, C*) and *ϕ*_*s*_ × *s*(*C*), respectively.

To establish a baseline, we first consider how changes in vital rates impact coral cover in undisturbed environments. In this case, equilibrium cover is more sensitive to changes in growth and mortality, and considerably less sensitive to changes in recruitment (Fig. 4A). Large (> 25%) decreases in growth eventually cause equilibrium cover to be lost at an accelerating rate, while non-linear effects of changes in mortality and recruitment are less pronounced. Next, we consider disturbed environments by simulating the same two disturbance regimes that we considered for Fig. 3. For each disturbance regime, we simulated 400 years of dynamics, and recorded average coral cover over the last 200 years. These simulations show that disturbance makes coral cover more sensitive to changes in any of the three vital rates (Fig. 4B,C). In other words, disturbance and changes in vital rates interact, as the effect of both combined is greater than the sum of the individual effects. In disturbed environments, average cover is still less sensitive to changes in recruitment than to changes in growth or mortality, but changes in any of these inputs have more pronounced effects under disturbed conditions.

**Figure 4:**
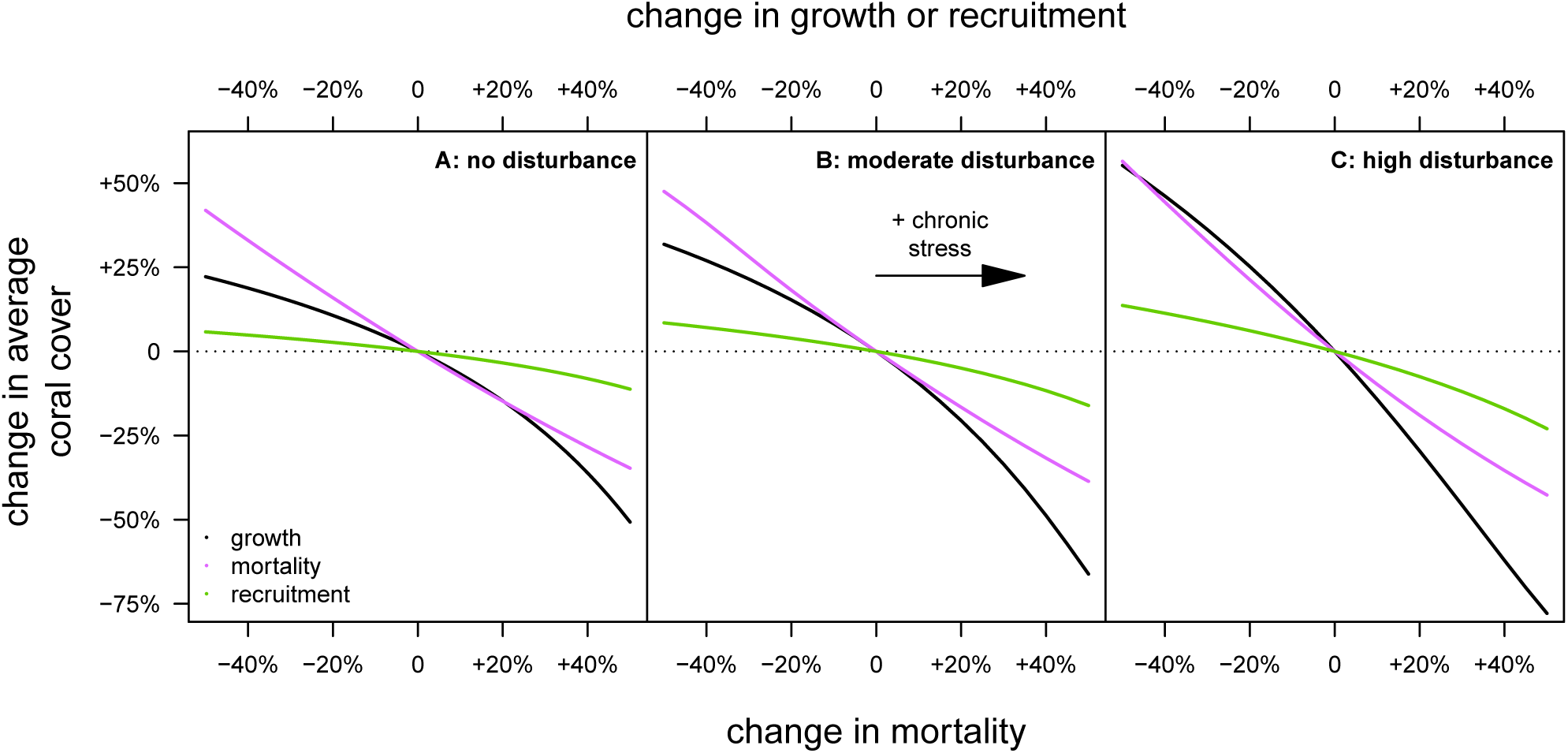
Average coral cover is more sensitive to changes in vital rates in disturbed environments. Each panel shows the proportional change in average coral cover in response to a proportional change in growth (black), mortality (violet), or recruitment (green). Within each panel, changes in vital rates are shown so that the putative effect of chronic environmental stress increases from left to right. A: No disturbance. B: Infrequent, moderate disturbance. C: Frequent, severe disturbance. Results in panel A are based on equilibrium calculations, while results in panels B and C are based on stochastic simulations.

To understand why small changes in vital rates have a bigger effect on coral cover in disturbed environments, we calculated the elasticity of total coral cover to changes in demographic inputs during the recovery period immediately following a population wipe-out (Fig. 5). Elasticities here are defined in the usual way, as the proportional derivatives of the relationship between coral cover and the demographic multiplier *ϕ* evaluated at *ϕ* = 1 (de Kroon et al., 1986). We compute the elasticity with respect to the direction of change caused by chronic environmental stressors (e.g., a decrease in growth or recruitment, and an increase in mortality). All elasticities are calculated with finite-difference approximations.

**Figure 5:**
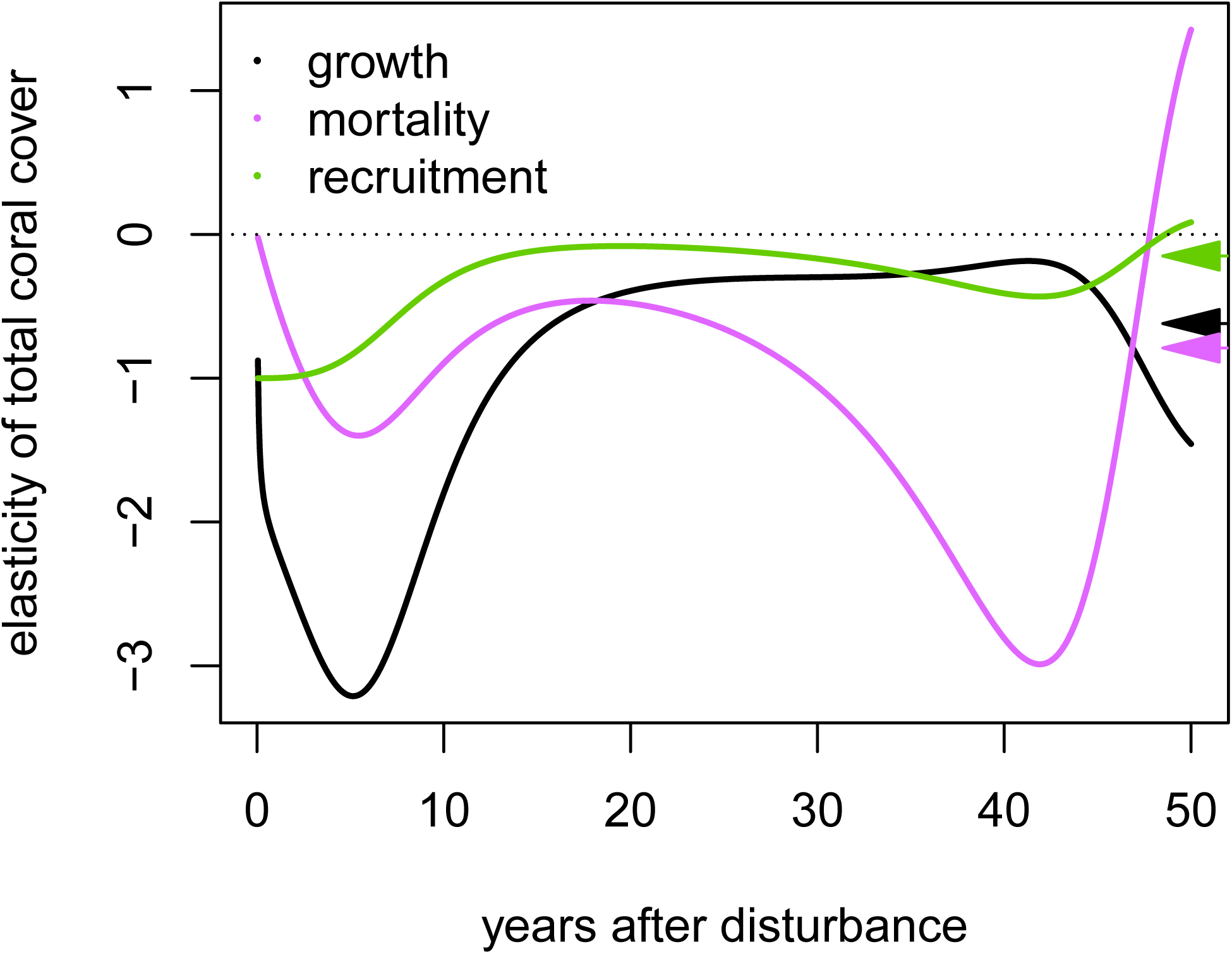
Elasticity of coral cover to vital rates changes considerably during the years following a population wipe-out. Lines show the elasticity of total coral cover to changes in colony growth (black), mortality (violet), and recruitment (green), as a population rebuilds following a population wipe-out. Elasticities are defined with respect to the change in vital rates caused by chronic environmental stress; that is, with respect to a decrease in growth or recruitment, and with respect to an increase in mortality. Arrowheads on the right axis show the long-run elasticity of the equilibrium cover in undisturbed environments.

In the immediate aftermath of a population wipe-out, coral cover is most sensitive to recruitment and growth of new colonies, and only minimally sensitive to mortality (Fig. 5). However, in the first few years of recovery, cover becomes increasingly sensitive to growth and mortality, and less so to recruitment. These elasticities are larger (in magnitude) than the elasticities observed at equilibrium (shown on the right axis of Fig. 5), especially for growth and recruitment. Coral populations are more sensitive to changes in vital rates after a disturbance because density-dependent crowding buffers the effect of demographic change in the long run, but acts only weakly in the uncrowded conditions created by a disturbance. As recovery continues, a more complicated pattern emerges, because changes in growth and mortality also affect the timing of the transient oscillations in population size. Despite these complicated patterns, cover remains only minimally sensitive to recruitment once the first few years of recovery have passed.

## Discussion

This analysis suggests several insights into the population dynamics of reef-building corals. First, size structure and space competition can drive surprising dynamics even in populations that routinely experience mass mortality events. Roughgarden et al. (1985) and others have already established that size structure and space-limited recruitment can lead to population oscillations in sessile marine invertebrates in undisturbed conditions; we show here that those oscillations appear in models with size-dependent growth and mortality schedules similar to those observed in a common coral species complex, *Pocillopora verrucosa*.

For many of today’s reefs, intermittent mortality pulses will disrupt the formation of regular population oscillations. However, recognizing that size structure causes coral cover to tend towards oscillation allows us to better project coral recovery in disturbed environments, for two reasons. First, when a new cohort of recruits settle and grow following a local extirpation (or near extirpation), coral cover can increase to levels that temporarily exceed the long-term carrying capacity. If a mortality pulse occurs at the peak of a nascent oscillation, coral cover should not necessarily be expected to rebound to pre-disturbance levels, even in a benign post-disturbance environment. Second, fluctuations in size structure explain variation in the recovery of coral populations immediately after a moderate disturbance. Recovery after a moderate disturbance is driven by the growth of surviving colonies, and thus cover will rebound most quickly if the surviving colonies are able to grow rapidly when intraspecific space competition is relaxed. In our model, mortality pulses acted independently of size, so size structure prior to a moderate disturbance dictated the pace of the post-disturbance rebound. If, instead, disturbance mortality depends on colony size, then the pace of recovery will be determined both by the size structure of the population at the time of the disturbance, and the size-selective mortality.

This analysis also suggests that occasional mass-mortality events make coral cover more sensitive to changes in vital rates driven by chronic environmental stressors. In the absence of disturbance, intraspecific space competition partially buffers the effect of changes in vital rates. Mass-mortality events continually reset coral populations to low densities where competition for space is less intense, and thus density-dependence is less effective at buffering simultaneous changes in underlying vital rates. Although we have presented our results as disturbance exacerbating coral sensitivity to chronic stressors, of course this interaction must flow in the other direction as well: chronic stressors will also make average coral cover more sensitive to changes in the frequency and magnitude of disturbance.

Finally, these results also suggest that coral populations will be more sensitive to changes in colony growth or mortality rates than to changes in recruitment, under all of the disturbance scenarios we have considered (Fig. 4). The relative robustness of coral cover to changes in recruitment is noteworthy because our model assumption of constant recruitment is a clear simplification. In reality, we expect external recruitment to vary through time, likely substantially (Graham et al., 2008; Thompson et al., 2018). However, because recruitment has only a mild impact on average coral cover, it follows that coral populations should be relatively insensitive to fluctuations in external recruitment. Of course, one population’s external recruits are another population’s spawned gametes, and thus region-wide declines in coral cover must trigger comparable declines in recruitment across a metapopulation, if gamete production in corals is proportional to total cover. The region-wide, metapopulation dynamics of several coral reefs coupled by larval migration, complete with local size structure and density-dependence, would be an intriguing topic for further study.

Our model makes a number of additional simplifying assumptions, any one of which provides scope for additional study. First, we have assumed that internal recruitment (that is, self-seeding by local colonies) is negligible. Empirical evidence for the importance of internal recruitment varies widely (Sammarco and Andrews, 1989; Gilmour et al., 2009; Jones et al., 2009; van Oppen et al., 2011). Additional simulations of our model (not shown) suggest that self-seeding dampens population oscillations, because high coral cover generates high larval production, which partially counteracts the reduction in recruitment caused by a dearth of available substratum. Second, we have assumed that when a coral colony dies, the space that it occupied immediately becomes available to living colonies or new recruits. This assumption is more appropriate for some causes of mortality, such as hydrodynamic dislodgement, than others, such as corallivory or bleaching. In these latter cases, dead coral skeletons need to be removed through breakage, bioerosion, or dissolution before surviving colonies or new recruits can occupy the newly available space. This lag between a colony’s death and the removal of its dead skeleton may further promote population oscillations in coral cover.

Third, we have assumed that changes in colony growth and mortality caused by chronic stressors act independently of colony size. For mortality at least, we expect that size-dependent mortality is driven by multiple mechanisms. Namely, when colonies are small, mortality is likely inversely related to size because smaller colonies are most susceptible to overgrowth by conspecific or heterospecific space competitors (Ferrari et al., 2012). On the other hand, when colonies are large, mortality is positively related to size because large colonies are more susceptible to dislodgement from hydrodynamic stress (Madin et al., 2014). If, for example, a chronic stressor such as ocean acidification increases mortality by making coral skeletons less dense and hence more brittle (Fantazzini et al., 2015), we might expect the mortality of large coral colonies to increase more rapidly than the mortality of small coral colonies. If data were available to quantify the size-dependent effects of chronic stressors more precisely, it would be straightforward to incorporate those effects into this modeling framework.

Finally, and in a separate, methodological vein, our model illustrates both the strengths and limitations of using physiologically structured population models (de Roos, 1997) to study coral populations. PSPMs can readily accommodate size-structure and density-dependence in a continuous-time framework that may be appropriate for the tropical environments that many coral reefs inhabit. As these results show, intraspecific density-dependence can be particularly important for elucidating how corals will respond to the combined effects of disturbance and chronic environmental stressors. On the other hand, PSPMs are less well suited to accommodating stochastic growth trajectories of individual colonies, which may also be important for gaining a more complete picture of coral population dynamics. Thus, PSPMs complement alternative approaches to modeling size-structured populations such as the population projection matrices (Hughes, 1984; Caswell, 2001) and integral projection models (Easterling et al., 2000; Madin et al., 2012), both of which are less well suited to handling density dependence, but which accommodate stochastic growth more readily.

## Acknowledgments

We thank the staff of University of California Gump Research Station for making our stays there enjoyable and productive. We gratefully acknowledge the support of the National Science Foundation: TEH and ASF were supported by award DMS 12-46991, PJE and RCC were supported by awards OCE 14-15268 and 16-37396, and KG was supported by award OCE 14-15300.

## Supplementary material

### Parameter values and justifications

#### *x*_0_, diameter of a newly settled coral spat

We use a value of 0.4 mm, which is loosely based on Babcock (1991).

#### *x*_max_, maximum diameter of a coral colony

Veron’s online factsheet reports that *P. verrucosa* colonies are “seldom more than 0.5 metres across” (http://www.coralsoftheworld.org/, accessed Jan. 20, 2020).

#### *g*(*x, C*), radial growth rate of a coral colony

To estimate the growth rate of *P. verrucosa*, we used annual monitoring data from the Mo’orea Coral Reef Long-Term Ecological Research (LTER) project for 2011 – 18. We use monitoring data from LTER sites 1 and 2 (both on Mo’orea’s north shore) and from 10 m depth on the outer reef. Monitoring consisted of annual visits to a permanent 40 m transect that contained several 0.5 m × 0.5 m quadrats spaced roughly 0.5 – 2 m apart (Edmunds, 2018). Quadrats were photographed in March or April of each year. Using the digital images, the diameter of the major and minor axis of each colony was recorded. The effective diameter of a colony is twice the geometric mean of the radii of the major and minor axis. To estimate the growth-rate function, we use pairs of data for which colonies can be (a) unequivocally identified as the same colony in consecutive years, (b) did not exhibit full or partial mortality (to the extent that observers could tell), and (c) remained entirely within the photographed quadrat in both years. Our analysis uses *n* = 1003 pairs of sizes in consecutive years available for analysis. Some of these pairs are generated by the same colony (for example, a single colony observed in four consecutive years generates three pairs of consecutive observations), and thus would be expected to generate correlated residuals. Thus, the effective sample size is somewhat smaller than 1003, and the actual standard errors associated with our fitted growth curve are larger than a naive calculation would suggest.

We assumed that the growth increments that we observe reflect density-dependent growth rates. In other words, we expect that observed growth was reduced by crowding. For each year in our time series, we calculated the total coral cover across all quadrats at 10m depth for each LTER site. We then calculated a density-independent growth increment by dividing the observed growth increment by the coral cover in the initial year. That is, if a colony is observed to have size *x*_*t*_ at year *t*, and size *x*_*t*+1_ at year *t* + 1, and if the average coral cover at the particular LTER site in year *t* is *C*_*t*_, then the density-independent growth increment is calculated as *y*_*t*_ = (*x*_*t*+1_ − *x*_*t*_)/(1 − *C*_*t*_). We then regressed *y*_*t*_ vs the size at the beginning of the time interval *x*_*t*_, using a quadratic regression that is constrained to generate a fitted value of 0 when *x*_*t*_ = *x*_max_ (Fig. S2). Our fitted equation takes the form *y*_*t*_ = *b*_0_ + *b*_1_ *x* + *b*_2_ *x*^2^ + *ϵ*, where the *ϵ*’s take the usual assumptions of independent and identically distributed Gaussian error. In other words, our fitted equation is

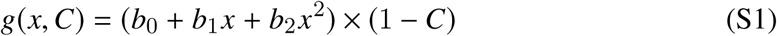

For our data, a constrained least-squares fit yields estimates of

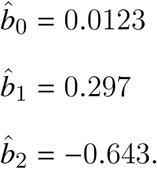

We do not report standard errors because accurate calculation of standard errors would require accounting for correlations among observations from the same colony, which is beyond the scope of this analysis.

Finally, note that by equating a growth rate *g*_0_(*x*) with an annual growth increment, we are essentially assuming that the growth rate is constant over the course of an entire year. Of course, in our model, the growth rate changes as the colony grows, so that the equivalence between the growth rate and the observed annual growth increment is inexact. However, growth rates are sufficiently slow, and change sufficiently slowly with changing size, that the loss in fidelity from using an observed annual growth increment to estimate a growth rate is minor.

#### *µ*(*x, C*), coral mortality rate

First, we assume that coral mortality is independent of coral cover. To estimate the size-dependence of coral mortality, we use data from corymbose corals on Lizard Island, Australia from Madin et al. (2014) (helpfully provided as a supplement to their paper). We modified their fit as follows. Because our model requires a mortality rate instead of an annual mortality, we fit a generalized linear model (GLM) with a complimentary-log-log (cloglog) link; Madin et al. used a more familiar logit link. Madin et al. used the log of the colony size as their predictor; however, fitting a quadratic model (on the link scale) with log diameter as the predictor yields very high mortality rates for small colonies (annual mortality rates of > 99.9% for colonies at *x* = 3 cm) that are not consistent with either our monitoring data or recent fits to similar data that appear in Kayal et al. (2018). Thus, we instead opted for a quadratic fit (again, on the link scale) with colony diameter as the lone predictor, even though this model is slightly AIC-worse than the fit with log diameter as the predictor (Δ AIC = 4.07). In terms of a mortality rate, this fit implies the following functional form for *µ*(*x, C*):

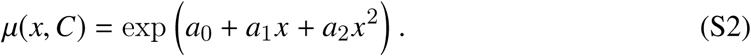

Holbrook et al. (2018) report that, on the north shore of Mo’orea, the annual mortality of newly detectable *P. verrucosa* colonies (which they define to be *x* ≈ 3 cm) is 35%. The fit Madin *et al*.’s data suggests an annual mortality for *x* = 3 cm corals of 61% per year. Thus we adjusted the *a*_0_ term in the model to give an annual mortality of 35% for colonies of size *x* = 3 cm. Our estimated, adjusted coefficients are

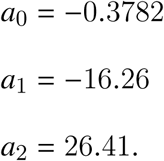

#### *s*_0_, coral recruitment rate

Holbrook et al. (2018) report that visible *P. verrucosa* recruits (2 ≤ *x* ≤ 3 cm) are found on the north shore of Mo’orea at an average rate of approximately 20 recruits m^−2^ y^−1^. Our growth and survival curves suggest that in uncrowded conditions roughly 40% of all recruits will survive long enough to become detectable, giving an arrival rate of newly settling recruits of roughly *s*_0_ ≈ 20/.4 = 50 recruits m^−2^ y^−1^.

### Computational details

The equilibrium of the undisturbed model (used to generate Fig. 4A and the stable stage structure shown in Fig. S1) was calculated using the numerical methods described in Kirkilionis et al. (2001). All other results were obtained by simulations using the Escalator Boxcar Train method described in de Roos (1988) and de Roos (1997).

### Additional figures

**Figure S1:**
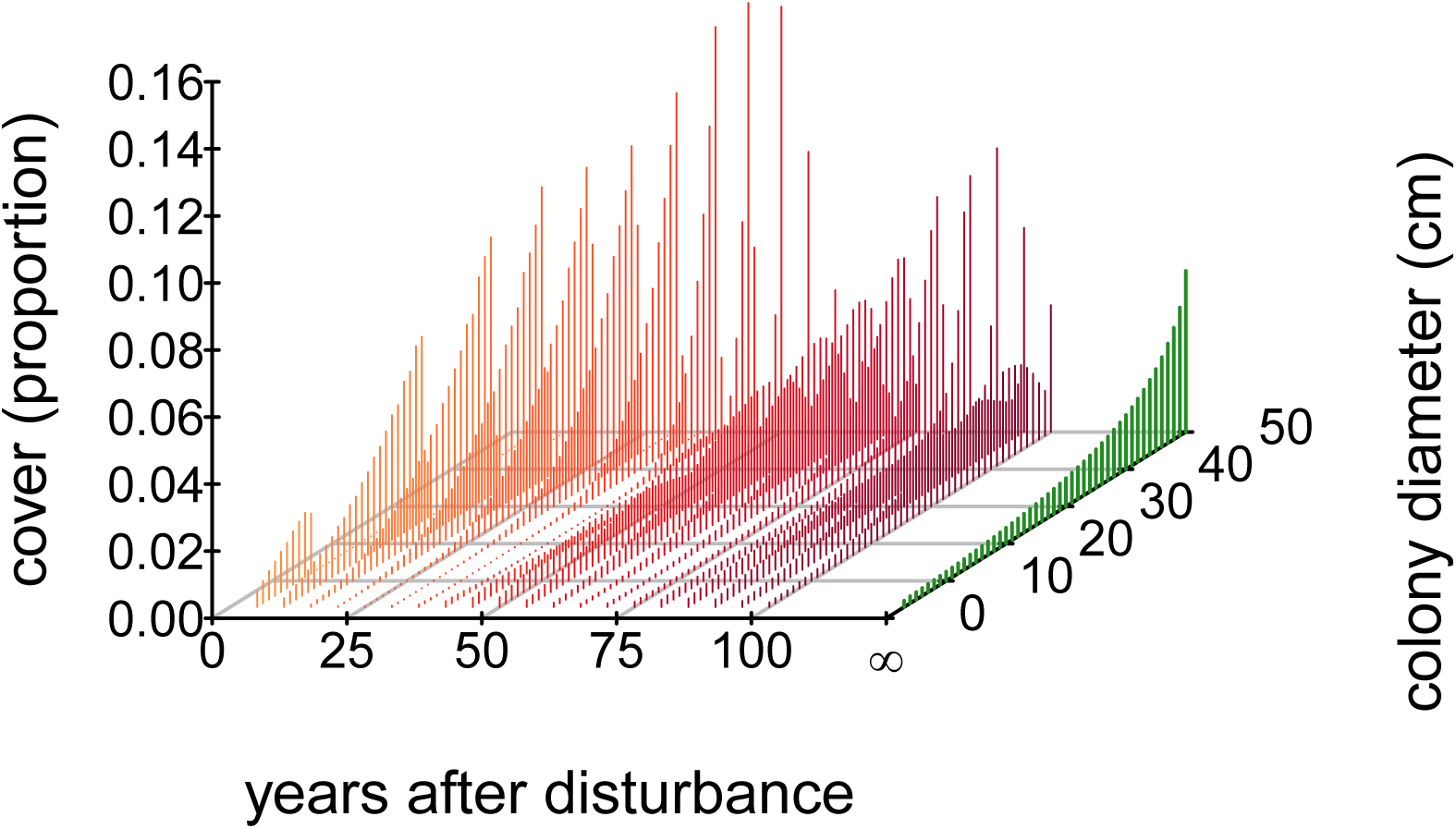
Oscillations in coral cover are driven by space competition between pulses of recruits. This display shows the size structure of the coral population for 100 undisturbed years following a catastrophe that completely eliminates existing coral cover. For ease of visualization, this display divides coral colonies into discrete size classes of 2 – 3 cm diameter, 3 – 4 cm diameter,…, 49 – 50 cm diameter, and shows the cover accounted for by colonies in each size class, expressed as a proportion of the total available space. Warm colors (orange and red) show size structure for years 5, 10, …, 100. The green histogram shows the equilibrium size structure to which the dynamics eventually converge.

**Figure S2:**
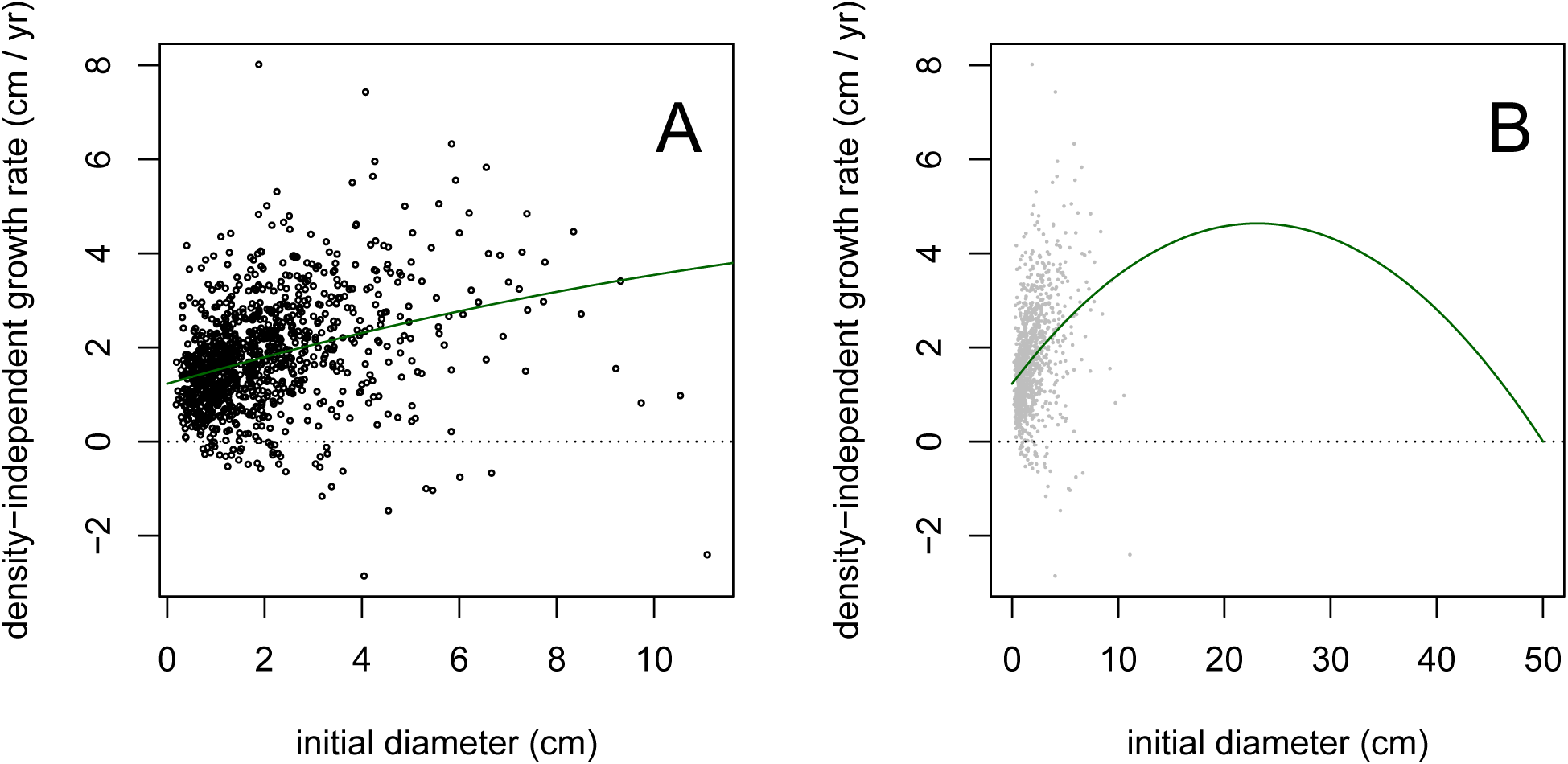
Estimated growth rate for *Pocillopora verrucosa*. A: Annual growth increment (corrected for crowding) vs. coral colony size for monitoring data from LTER sites 1 and 2 on Mo’orea’s north shore, and best constrained least-squares fit of eq. S1. B: As in panel A, but extended over the full range of coral colony sizes.

